# RosettaGPCR: Multiple Template Homology Modeling of GPCRs with Rosetta

**DOI:** 10.1101/2019.12.13.875237

**Authors:** Brian Joseph Bender, Brennica Marlow, Jens Meiler

## Abstract

G-protein coupled receptors (GPCRs) represent a significant target class for pharmaceutical therapies. However, to date, only about 10% of druggable GPCRs have had their structures characterized at atomic resolution. Further, because of the flexibility of GPCRs, alternative conformations remain to be modeled, even after an experimental structure is available. Thus, computational modeling of GPCRs is a crucial component for understanding biological function and to aid development of new therapeutics. Previous single- and multi-template homology modeling protocols in Rosetta often generated non-native-like conformations of transmembrane α-helices and/or extracellular loops. Here we present a new Rosetta protocol for modeling GPCRs that is improved in two critical ways: Firstly, it uses a blended sequence- and structure-based alignment that now accounts for structure conservation in extracellular loops. Secondly, by merging multiple template structures into one comparative model, the best possible template for every region of a target GPCR can be used expanding the conformational space sampled in a meaningful way. This new method allows for accurate modeling of receptors using templates as low as 20% sequence identity, which accounts for nearly the entire druggable space of GPCRs. A model database of all non-odorant GPCRs is made available at www.rosettagpcr.org.

**Author Summary:** Structure-based drug discovery is among the new technologies driving the development of next generation therapeutics. Inherent to this process is the availability of a protein structure for virtual screening. The most heavily drugged protein family, G-protein coupled receptors (GPCRs), however suffers from a lack of experimental structures that could hinder drug development. Technical challenges prevent the determination of every protein structure, so we turn to computational modeling to predict the structures of the remaining proteins. Again, traditional techniques fail due to the high divergence of this family. Here, we build on available methods specifically for the challenge of modeling GPCRs. This new method outperforms other methods and allows for the ability to accurately model nearly 90% of the entire GPCR family. We therefore generate a model database of all GPCRs (www.rosettagpcr.org) for use in future drug development.

## Introduction

### G-protein Coupled Receptors Represent Important Therapeutic Targets

G-protein coupled receptors (GPCRs) are the largest family of membrane proteins in the human body comprising nearly 800 distinct receptors [1]. They orchestrate cellular response to extracellular signals and thus play roles in immune response, cardiopathies, and neural development. They are a ubiquitous family of proteins evolved over time to respond to a variety of stimuli including ions, small molecules, larger peptides, and even light [2]. Given their position at the interface of a cell with its environment, they are attractive targets for therapeutic intervention. Current estimates suggest around 30% of drugs available act at a GPCR [3].

### Experimental structures of GPCRs are determined at an increasing rate overcoming substantial obstacles

The first atomic resolution structure of a GPCR was rhodopsin in 2000, in part due to its high abundance and stability from native sources [4]. For most receptors, expression levels are well below what is needed for structural characterization from orthologous sources. Therefore, it wasn’t until 2007 that the structure of a second receptor was experimentally determined [5,6]. As dynamic, membrane-bound proteins, significant protein engineering was needed for structure determination (i.e. thermostabilization through mutation, nanobodies, fusion protein, or truncation of flexible termini) [7]. Since 2007, about 50 unique receptor structures have been determined. While this is a tremendous achievement, this represents only about 6% of the GPCR superfamily. Even when focusing on non-olfactory GPCRs that are generally considered as druggable targets, nearly 350 unique receptors remain to be structurally characterized either for better understanding of how current drugs bind their targets or for structure-based drug discovery. Of importance, at least 100 of non-olfactory GPCRs have been designated orphan receptors due to a lack of chemical matter [8]. Knowledge of the structural details of the ligand binding pocket could assist in identifying chemical probes for these dark receptors.

### Computational Modeling can Extend our Current Understanding of GPCR Structures

Given this knowledge gap, homology modeling is an important tool for generating models of as-of-yet undetermined receptor structures. Homology modeling uses a protein template with a shared topology to map the target sequence onto its backbone coordinates in a process called threading [9]. Early homology modeling relied on a single template structure for target structure prediction. However, these methods fail to generate accurate models using templates with low sequence identity to the target protein. More recently, the use of multiple templates has seen success in modeling targets in which the sequence identity is below 50% to any given template [10,11]. Given that GPCRs share identities in the range of 20-30%, GPCR-specific homology modeling has largely moved towards multiple template modeling. Servers for the prediction of GPCRs from multiple templates are available including GPCR-ModSim [12], GPCR-I-Tasser [13], GPCRM [14], GPCR-SSFE [15], and GPCRdb [16]. GoMoDo, another server, uses single-template modeling [17]. The underlying software for all these servers is Modeller [18], except for GPCR-I-Tasser [11]. To date, no GPCR-specific multi-template modeling method has been developed in Rosetta, a software capable of structure prediction, design, and docking of ligands [19,20]. Despite a unified method for GPCR modeling, Rosetta’s performance on single-template modeling of GPCRs has been analyzed in the past with mixed success. In the GPCR Dock experiment [21], Rosetta performed best in the structure prediction of the Smoothened receptor ligand binding pose [22].

### Rosetta Hybridizes Multiple Templates

While other methods predefine template segments for various parts of the target model or averages template structures, Rosetta handles multiple templates simultaneously during its modeling process [23]. Rosetta holds all templates in a defined global geometry and randomly swaps parts of each template using Monte Carlo sampling to identify regions from the various templates that best satisfy the local sequence requirements. This template swapping occurs in parallel with the traditional peptide fragment swapping from a database derived from the PDB based on the target sequence and predicted secondary structure, a hallmark of Rosetta’s folding algorithm [24]. This simultaneous sampling of template segments and peptide fragments allows the energy function to define which segments to keep from the various templates based on how well each segment improves the overall score of the model. Hybridization of templates has been shown to be successful in CASP experiments, particularly for low template identity targets down to 40% [23]. Below 40% identity, Rosetta is capable of producing accurate models, though it is not known *a priori* if the output models will be reliable.

### Development of a GPCR-specific Multiple-Template Homology Modeling Protocol in Rosetta

Given the past success of Rosetta in single template homology modeling of GPCRs [22] and the novel strategy of multiple template modeling in the Rosetta framework [23], we set out to develop a protocol specific to GPCRs that utilized these new algorithms. The change from the previous single-template homology modeling to multiple-template modeling was multifaceted and we tested each component individually. First, the use of multiple templates begs the question of what is the optimal number of templates to use. Previous work in multiple template homology modeling suggested that there is a goldilocks effect in which multiple templates are better than one but too many templates could actually hurt the modeling process [25]. Additionally, as Rosetta uses a peptide fragment library, we evaluate its influence on enhancing modeling accuracy. Further, loop closure is handled simultaneously in Rosetta’s multiple-template homology modeling through the use of these peptide fragments. As these loops are defined by their input template, we optimized the alignment in these regions and tested its effect on model accuracy. We benchmarked our methods on 34 available structures of unique GPCRs covering the four classes (A, B, C, and F) that had structures available at the outset of the study in July 2017. Additionally, we chose to model all targets using templates below 40% sequence identity, unless otherwise noted, to mimic the situation when predicting novel target structures. We find that our GPCR-specific Rosetta-based multiple template homology modeling method (RosettaGPCR) is highly accurate due to the curated sequence alignments and peptide fragment utilization. We find, that our new method accurately models Class A receptors down to a template identity of 20%. Further, this method outperformed other GPCR servers in the prediction for four new GPCR structures. Based on this success, we established a database of all human non-odorant receptors available for use (www.rosettagpcr.org). Altogether, RosettaGPCR is currently one of the best available methods for modeling this pharmacologically important family of proteins.

## Methods

### Description of Benchmark Data Set

For this study we chose to model 34 crystal structures of GPCRs covering the four structurally characterized classes A, B, C, and F. In total there were 29 Class A members, two Class B receptors, two Class C receptors, and one Class F receptor (Table S1). Importantly, we chose to model the receptors using exclusively templates below 40% sequence identity, unless explicitly noted, as this most closely resembles the majority of real-life cases when modeling GPCRs.

### Generation of multiple-sequence alignments of templates

Initial alignments for the benchmark set were obtained from the GPCRdb [26]. This largely ensured that the transmembrane α-helices were well aligned. To improve on these alignments, the structures were visualized in PyMol, and the structural alignments were compared to the sequence alignments. Transmembrane helical sequences were aligned starting from the most conserved residue in each α-helix and extended outwards using the structural alignments to guide insertion and deletions along the α-helical axis. Loop alignments were generated based on the alignment of vectors of Cα to Cβ atoms between receptor structures. If structures were present in loop regions such as disulfides, α-helices, or β-sheets, these were preserved in the alignment. Remaining residues that could not be aligned by any of the above metrics were moved to be adjacent to a region of defined secondary structure to ensure proper fitting of peptide fragments between ordered and unordered regions. The alignment of the 34 receptors is shown in Figure S1. Additional alignments were generated using the default options of ClustalOmega [27], Muscle [28], T-Coffee TM-PSI and Espresso [29], and Mustang [30] and used without further modification.

### Template Selection

For all receptors, a pairwise identity matrix was generated using ClustalOmega [27]. The reported identities were used to rank the templates for each receptor model. Shown in Table S2 is the ranked list of templates for each target receptor. While most templates have sequence identities below 40%, those highlighted in yellow were removed because they featured sequence identity above the 40% threshold. Templates labeled in bold were used in single-template high identity modeling to compare to previous benchmark [22].

### Generation of Additional Input Files

Membrane spanning topology files were generated by submitting the sequence of the target proteins to Octopus [31]. The output files were converted into Rosetta readable span files with Rosetta’s built in octopus2span.pl script. Disulfide bond restraint files were prepared for each target protein for the conserved disulfide bond between TM3 and ECL2, except for LPA1 and S1P1. Additional disulfide bonds within ECL3 were mapped as needed.

### Sequence alignment of target sequence to template MSA

Alignment of sequences without known structure was accomplished similarly as above. First, alignments were extracted from GPCRdb. Then the highly conserved residues in each helix were aligned with the template MSA. Positioning of residue x.50 (BW numbering) often corrected helix alignments but gaps and deletions were propagated throughout families. For receptors lacking the highest conserved residue in each helix, other motifs such as DRY, NPxxY, and CxxP were used for helix positioning. The loops were aligned as such: ECL1, ICL1, and ICL2 were aligned using common sequence motifs (eg. xWxxG in ECL1). For ICL3, sequence alignments were maintained within families particularly with receptors with short loops such as the Class B and chemokine receptors. For the majority of receptors, the ICL3 sequence was split at the halfway point between TM5 and TM6 and the halfves were adjoined directly to the end of TM5 or beginning of TM6, respectively. For ECL3, particular attention was given to the presence of cysteines for either internal ECL3 disulfides or disulfides between N-terminus and ECL3. These cysteine residues were used for alignment. For receptors lacking cysteine bonds, patterns identified in template familes were used to fix family alignments. Remaining receptors, again had the sequence of the loop halved and adjoined the halves to their next helix sequence. For ECL2, targets were grouped by putative ligand binding type: aminergic, lipid, peptide, unknown. Based on this grouping, the alignments were carried out specifically for their family type (i.e. a beta sheet was predicted and aligned for all peptide receptors). For receptors with unknown ligand type and dissimilar ligand, the loop sequence was first divided at the conserved cysteine residue and this residue was aligned generating two shorter loops. These loops were then halved and adjoined to their nearest fixed structural feature, either a transmembrane helix or the conserved disulfide bond with TM 3. The full MSA of all receptors is available at www.rosettagpcr.org.

### Model Production

With all input files in hand, target sequences were threaded onto the pre-aligned templates using Rosetta’s partial_thread application [23]. Threaded models were passed to the hybridization application via use of Rosetta XML scripts [23,32]. Either 100 models or 1000 models were generated per run as noted in the text.

### Data Availability

Top models for each receptor are available at www.rosettagpcr.org. All scripts and files needed for generating the models are also provided as a protocol capture on this website.

## Results

### Blended Sequence- and Structure-Based Alignment Is Critical for Modeling Success

Inherent to any homology modeling protocol is an alignment between the sequence of the target protein and the template structure. As different families of GPCRs share low sequence identity with one another, sequence alignment is not trivial for this class of proteins. The best-known alignment of GPCRs is Ballesteros-Weinstein (BW) numbering [33] which identifies the most conserved residue in each α-helix and sets as a starting point for alignment (#TM-span.50). Counting along the α-helix in reference to this residue all other residues are labeled. While highly useful, this alignment falls short in two areas. As receptor structures became available, it was found that not all receptor families adhere strictly to the *i* to *i+4* periodicity in every α-helix [34]. Insertions and deletions have resulted in local alterations of the helicity, in particular around proline or glycine residues, and thus the BW numbering, of certain subfamilies of proteins. Secondly, BW numbering fails in the loop regions as different receptors have varying lengths of α-helices and dramatically different loop structures within extracellular loop 2 (ECL2) adopting disordered regions, α-helices, or β-sheets. Further, as these loops are critical for ligand recognition [35], they diverge widely in sequence proving even more challenging for the creation of meaningful sequence alignments. However, despite sequence divergence, there is evidence for structural conservation in these regions [36]. Therefore, a critical component to the present method has been to blend sequence and structure information into an optimized knowledge-based sequence alignment for GPCRs (Fig S1). We compared this new alignment to other well-known sequence- or structure-based alignment methods. For each receptor in the benchmark, 100 models were generated for each of the six alignment methods tested. The average RMSD for a target protein was divided by the average RMSD for the same target using the new alignment resulting in a fold change and the average across the full benchmark is reported (Figure 1). As seen, despite using sequence-only (ClustalOmega [27] and Muscle [28]), structure-only (Mustang [30]), or automated blended alignments (T-Coffee, TM-PSI, and Espresso (PDB Mode) [29]), the knowledge-based alignment performs the best in all regions tested. For Class A receptors it is found that the transmembrane (TM) region is modeled nearly equivalently across all methods with the most improvement coming from improvements in ECL2 modeling. For Classes B, C, and F there is a large improvement in modeling of all regions, except for Mustang alignments of the TM region, the only metric and class in which Mustang outperforms our alignment. Importantly, for all classes, the accuracy of ECL2 is strongly improved demonstrating this new alignment to be critical for accurate modeling of this region.

**Figure 1:**
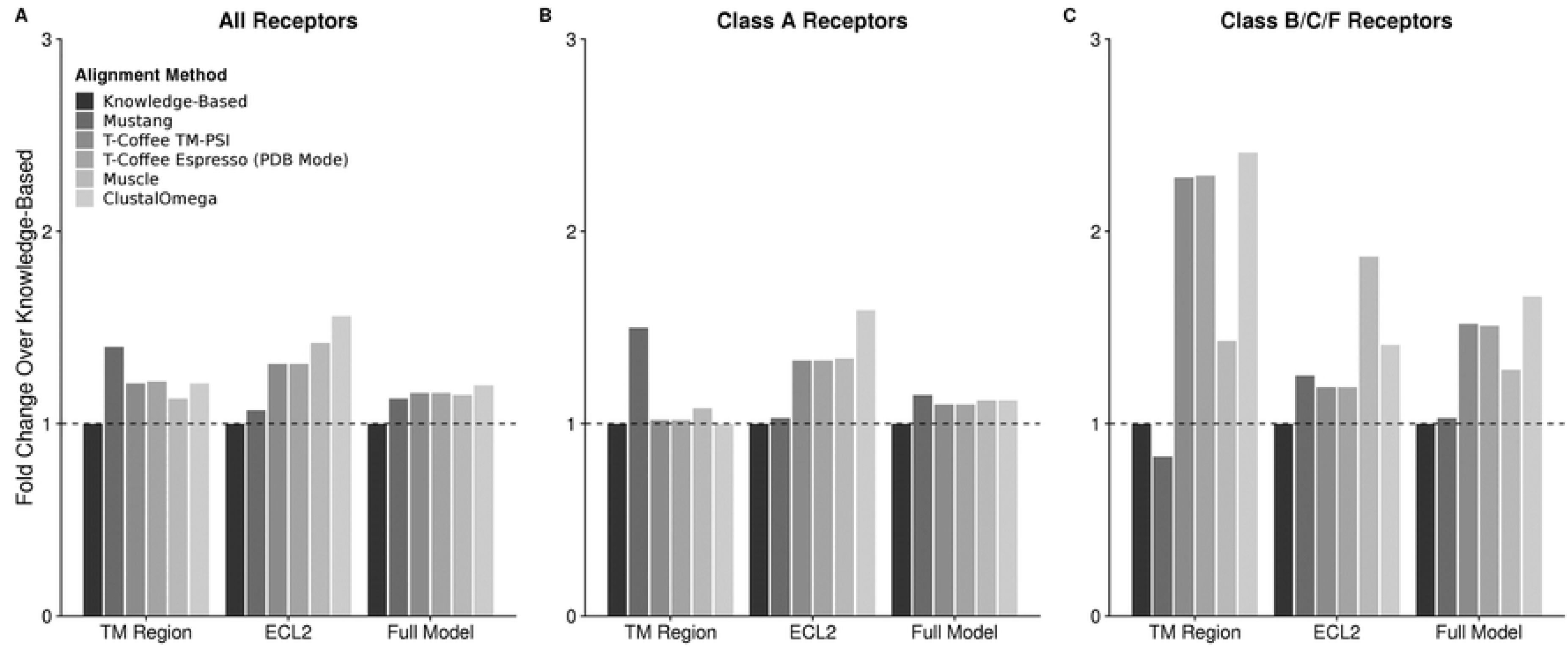
Comparison of Average RMSD Change using Various Alignment Methods. A total of 100 models were produced for each alignment method. The average RMSD of the models were normalized to the average RMSD of the models produced with the knowledge-based alignment (black). Values above 1 represent an alignment method that produced on average worse models while values below 1 represent an alignment method that produced on average better models. For (A) all receptors regardless of family, the knowledge-based modeling performs the best regardless of region analyzed. When split between (B) Class A and (C) Classes B, C, and F, the majority of the improvements are found the Classes B, C, and F where template availability is limited.

### Peptide Fragment Hybridization Improves Target Model Quality

Our previous benchmark of GPCR modeling relied on single-template threading [22]. We wanted to recapitulate this initial study using the Hybridize code [23] to allow for peptide fragment insertion but not template swapping to see what effect peptide sampling alone had on output quality. This benchmark dataset was limited to the eight receptors with high identity templates available in 2013 (β1AR, β2AR, M2R, M3R, δOR, κOR, μOR, and NOPFQ). In this experiment, each target was modeled on the single best available template with sequence identity either greater than 40% or less than 40% and allowed to hybridize with the peptide fragment library (Table 2). As seen in Figure 2, using the exact same template as was used in the previous threading-alone method, hybridization of template structures with peptide fragments can substantially improve output model accuracy in all measured regions. The transmembrane region improves on average by more than one Angstrom to 0.8 Å root-mean-square deviation (RMSD) to the crystal structures showing highly accurate modeling of this region. The ECL2 region also showed a dramatic improvement with an average RMSD to the crystal structures of 1.0 Å compared to the previous method of single-template modeling without peptide fragment insertion which reported an average RMSD of 5.0 Å. The full model RMSD, which accounts for all remaining loops and flexible termini, also showed modest improvement from 2.9 to 2.1 Å. These results were similar when using a single template with sequence identity less than 40%. Both the TM region and ECL2 improved by at least 1.0 Å while the Full Model RMSD actually worsened by 0.5 Å. Taken together, peptide insertion accounts for a substantial improvement over threading alone even when templates of sequence identity below 40% are used.

**Figure 2:**
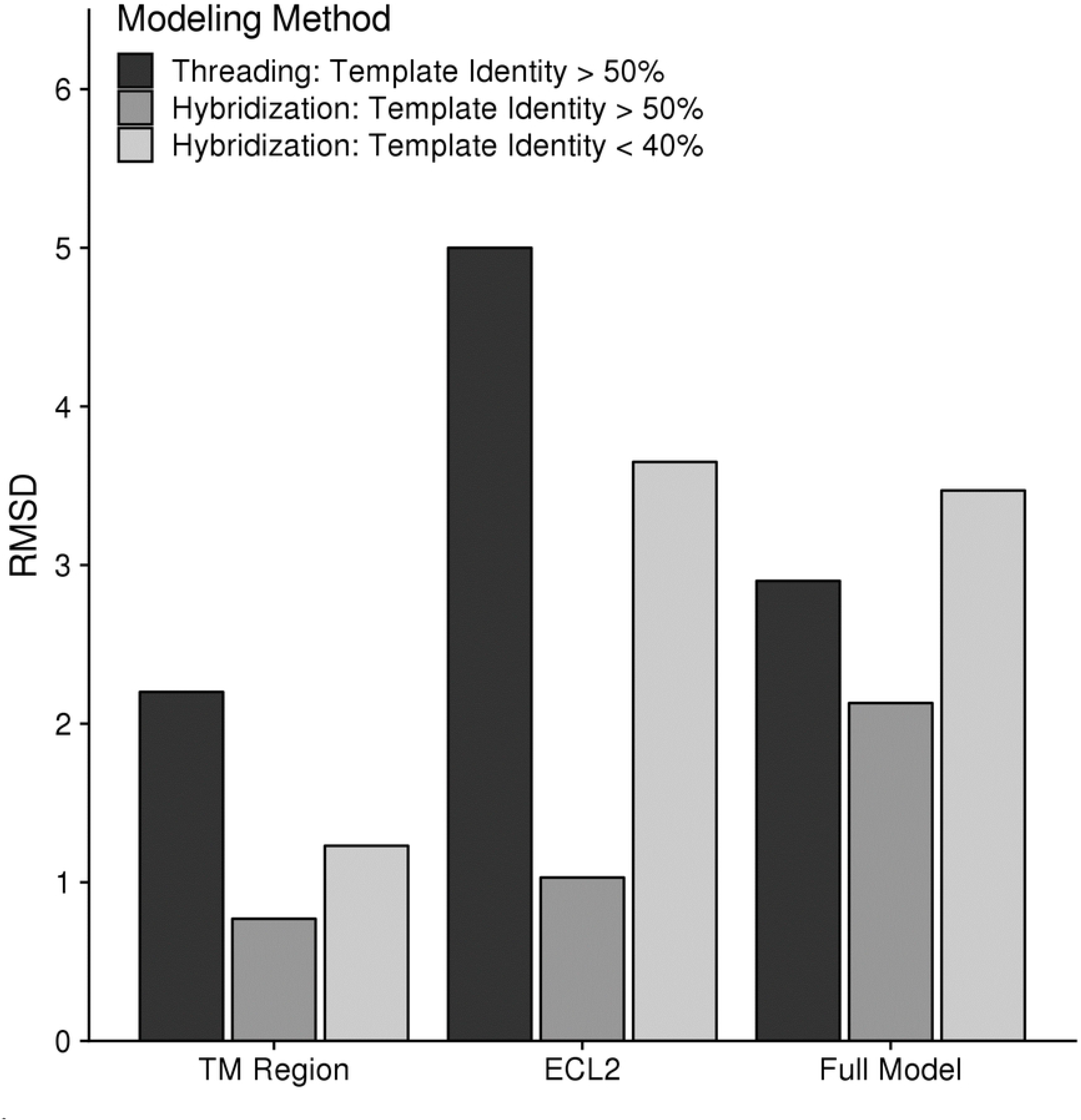
Comparison of Single Template Modeling Methods with Peptide Insertion. Using only a subset of receptors and templates that were available in our original GPCR modeling benchmark (yellow in Table S2), 100 models were generated using either a single high identity template or the best template available below 40%. Original results from the 2013 benchmark [22] are displayed in black. Using the hybridize code with the same original templates dramatically improved the results across all measures (medium grey). Using a low identity template in hybridize (light grey) expectedly worsened the results compared to the high identity template but was either better or comparable with the original threading alone algorithm.

### Multiple Templates Improve Performance for Low Sequence Identity Targets

While peptide insertion helped improved accuracy in the TM and ECL2 regions, overall model accuracy weakened when using a single template with sequence identity less than 40% to the target model. Therefore, we expected that multiple templates could overcome the shortcomings of any single template when modeling a target with low identity templates [23]. We generated 1000 models for every receptor using either the single best template less than 40% identity or the ten best available templates under 40% identity and compared the average RMSD of the resulting models (Figure 3 and 4). As expected, the average RMSDs improved for almost all receptors in the ECL2 and Full Model criteria. The TM region was rather insensitive to the increase in template availability showing on average only 0.05 Å improvement for the whole set. This is likely due to the high degree of structural similarity of the fold of inactive GPCRs. A few exceptions to the overall trends deserve attention. In the TM region, both class C receptors perform extremely poorly in either set of templates. This is due to the fact that the 40% threshold for selecting templates removed the other class C receptor from the template pool such that they were modeled with non-class C templates. As the structure of the TM region is distinct for these proteins compared to the other classes, the error was expected to be high. For the class B receptors, the two structures have a sequence identity of 35% with respect to one another allowing these structures to be included as templates in the benchmark. Therefore, the single template TM RMSD outperforms the ten template TM RMSD by nearly 0.5 Å. In ECL2, there are two class A receptors (S1P1 and LPA1) that perform extremely well when using a single template as compared to ten templates. These are the only two receptors in the benchmark that lack the conserved disulfide between ECL2 and TM3. Their loop structures are quite distinct from all other receptors and as a result, loop modeling only performs well when using the other as a template (Figure 4C). In the full model RMSD, both Rhodopsin and the Smoothened receptor perform extremely poorly regardless of the modeling method used. This is because both have extremely long and unusual loops and termini (Figure 4F). Of note, the TM regions of these two receptors are accurate with 1.5 Å and 2.7 Å RMSD to the crystal structure of 1U19 and 4JKV, respectively. Additionally, only two Class A receptors perform worse in the Full Model RMSD calculation when using multiple template. These again are S1P1 and LPA1 which performed poorly in the ECL2 modeling. It appears that the poor quality of ECL2 is reflected in the Full Model RMSD as the difference in the TM region for these two structures is only 0.1-0.2 Å.

**Figure 3:**
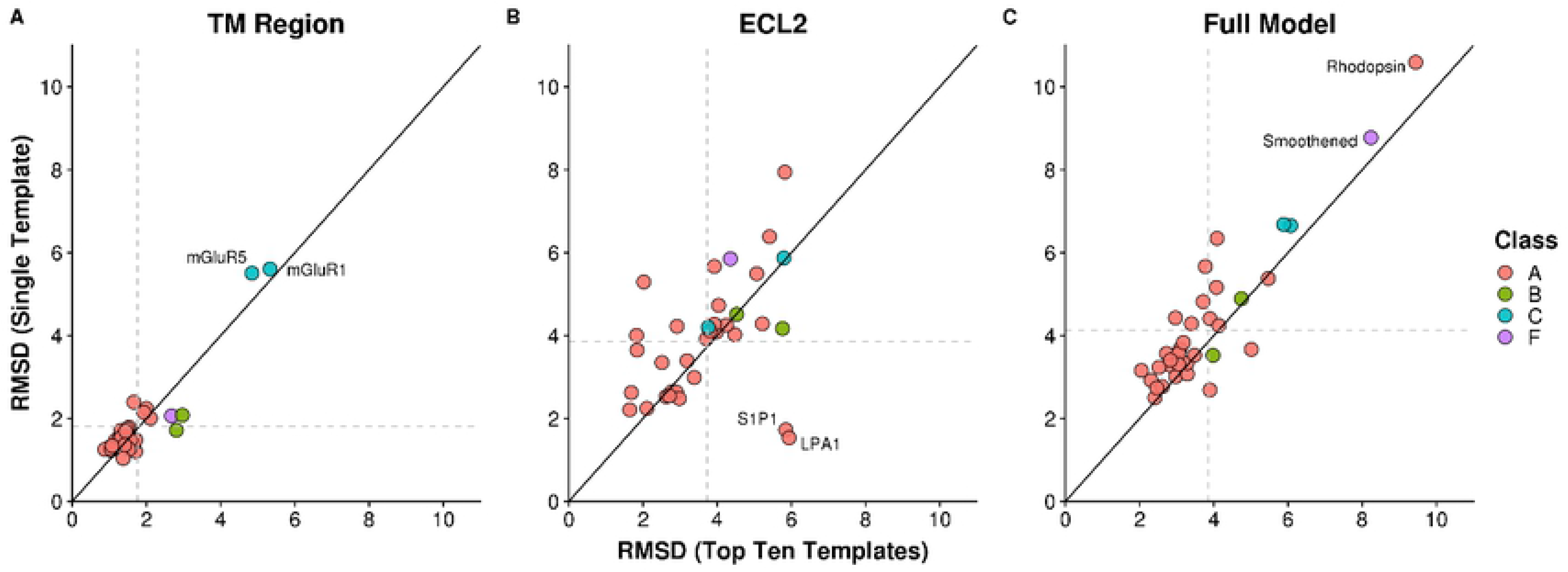
Comparison of Average RMSDs for Single versus Multiple Template Homology Modeling. Using either one template or ten templates, 1000 models were generated for each target and the average RMSD was calculated over the TM region (A), ECL2 (B), and the full model (C). Values that fall above the diagonal performed better when using multiple templates and values that fall below the diagonal performed better with a single template. Targets are colored by class.

**Figure 4:**
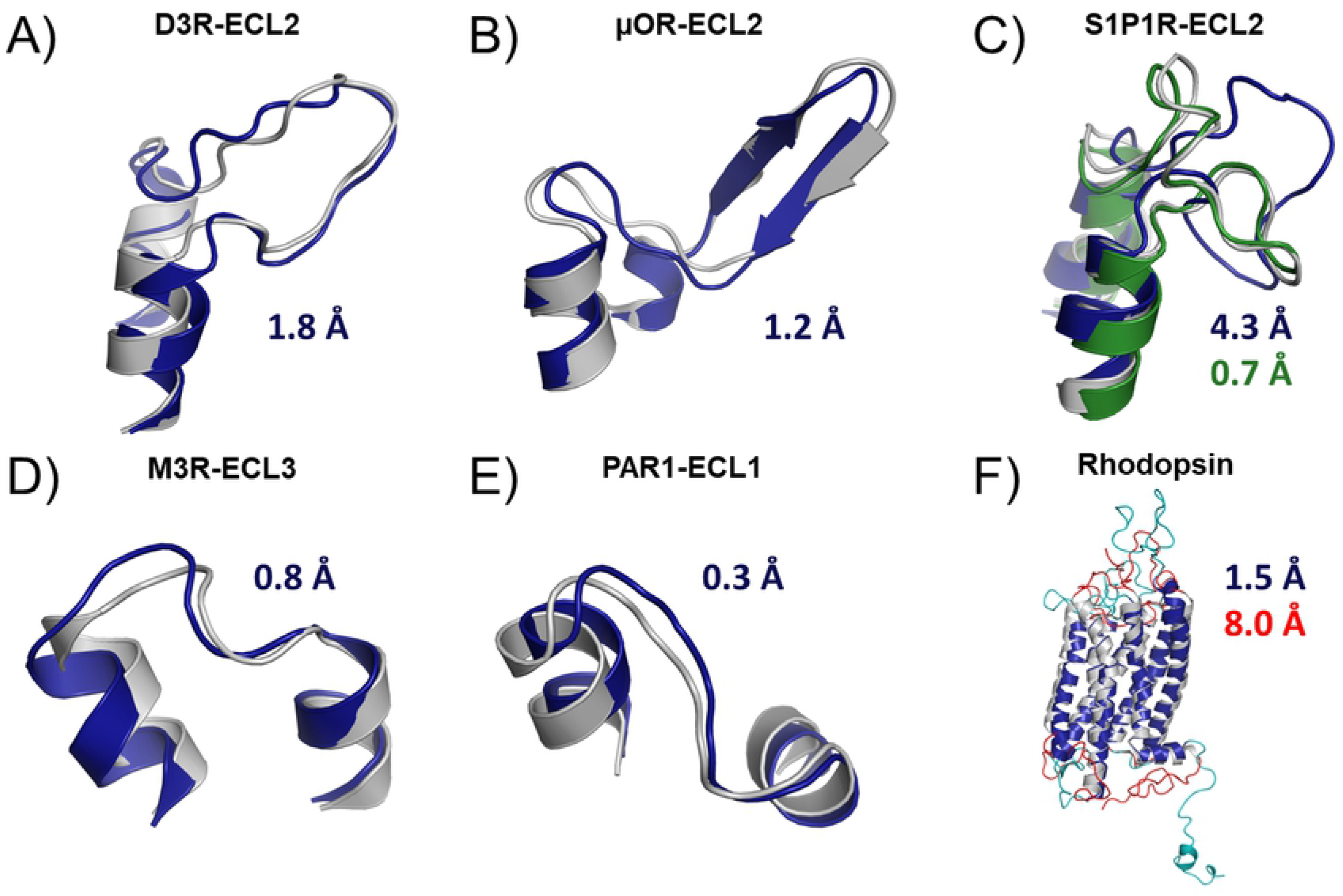
Examples of results obtainable with RosettaGPCR. In all cases, the crystal structure is colored grey and the model is blue. Three different ECL2 loops structures, unordered (A), β-sheet (B), and lipid (C). RosettaGPCR performs well on loops containing the conserved disulfide. For lipid receptors lacking the conserved disulfide (C) multiple templates (blue) perform worse than using a single template with similar structure (green), in this case the LPA receptor. Extracellular loops 3 (D) and 1 (E) also perform quite well with this method. In general, RosettaGPCR can model the TM region of most receptors below 2 Å (F). However, for receptors like rhodopsin with complex loop structures and termini (red), the model (cyan) fails to capture the overall conformation (8.0 Å RMSD).

### Identification of the Optimal Number of Templates

As reported previously for multiple template homology modeling, the use of multiple templates, while improving results over the single template approach, will weaken model accuracy if too many templates are used [25]. Thus, we determined an optimal number of templates for GPCR modeling using our method. We generated an additional 1000 models for each receptor using either five or all available templates and compared the data with the previous data on one and ten templates (Figure 5). For both the TM region and ECL2, using all available templates was worse than any other set of templates while the average RMSD were quite similar for one, five, and ten. However, for the full model accuracy, using a single template was worse than all other template sets, though comparably poor to the all available template compared to five or ten templates. Five templates performed distinctly well in the full model accuracy compared to the other template sets while only providing modest improvement over the other template sets in the TM and ELC2 regions. Therefore, we suggest five templates to be the best number of templates for modeling GPCRs with our method.

**Figure 5:**
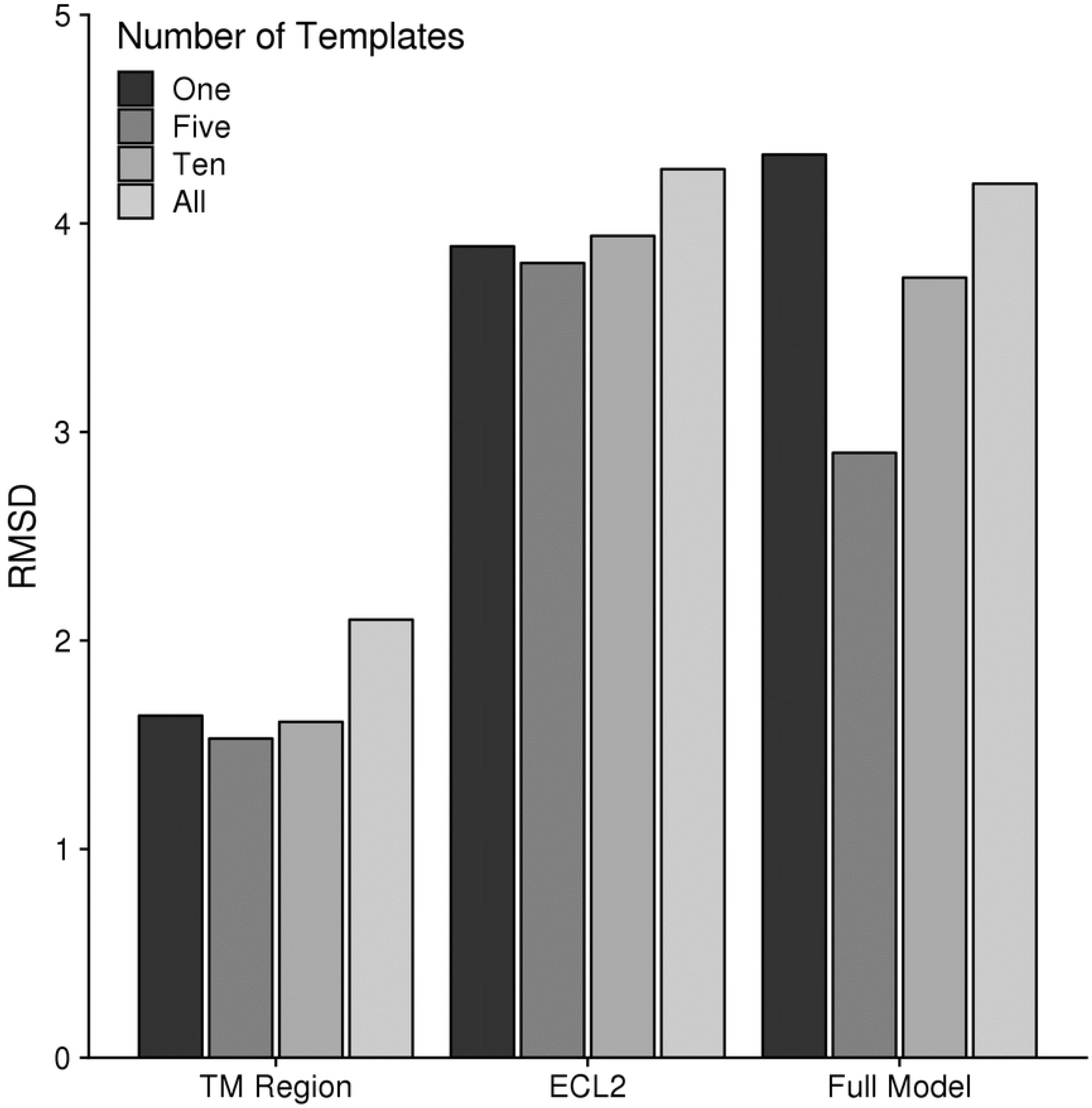
Comparison of Model Accuracy using Various Numbers of Starting Templates. For each target, 1000 models were generated using either 1, 5, 10, or all available templates. The average RMSD is plotted for the TM region, ECL2, and the Full Model.

### RosettaGPCR Outperforms other GPCR Modeling Servers

We next compared our method with other GPCR modeling servers. Three servers with publicly available databases of GPCR models, GPCRdb [16], GPCR-I-Tasser [13], and GPCR-SSFE [15], were included. Additionally, we utilized the GPCR-ModSim [12] server as it is very user-friendly and fast. We identified four human GPCR structures that were released following the conclusion of method development. The four structures were C5aR1, Y1R, PTAFR, and D2R (PDB IDs 6C1R [37], 5ZBQ [38], 5ZKQ [39], and 6CM4 [40], respectively). Of note, GPCR-ModSim, which is the only on-demand server we tested, had been updated to include the structures of PTAFR and D2R. GPCR-SSFE had the structure of D2R in its database. Therefore, we excluded results from these servers for these receptors. We generated 100 models of each receptor target using five template structures and selected the best model by total energy. In comparing our model with the models generated by other servers, we find that RosettaGPCR consistently outperforms the other approaches (Figure 6). Only one server performed on one target better than RosettaGPCR. GPCRdb had a better ECL2 of Y1R compared to ours with RMSDs of 1.6 Å versus 1.9 Å, respectively.

**Figure 6:**
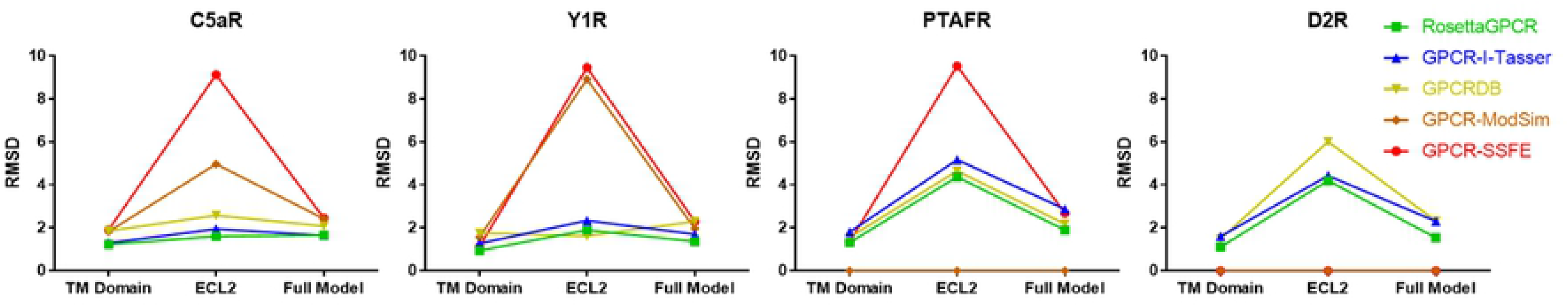
Results of Novel Structure Prediction from Various GPCR Modeling Servers. Blind predictions were carried out on C5aR1 (PDB ID 6C1R [37]), Y1R (PDB ID 5ZBQ [38]), PTAFR (PDB ID 5ZKQ [39]), and D2R (PDB ID 6CM4 [40]). 100 models were generated for each target with RosettaGPCR (black) and the best scoring model was used for analysis. The RMSD of each model for the various servers were calculated for the TM region, ECL2, and the full model. No data is available for GPCR-ModSim for D2R and PTAFR and for GPCR-SSFE for D2R because these servers had already included these targets in their database.

### Accuracy of Models with Increasingly Worse Templates Does Not Decline Linearly

In our previous work on GPCR modeling using single-template threading, it was found that templates needed to be greater than 50% sequence identity for accurate models [22]. Subsequently, the use of multiple-template modeling in Rosetta was suggested to be accurate to about 40% sequence identity [23]. However, in this current benchmark we only use templates with less than 40% sequence identity and still produce highly accurate receptor models. Therefore, we wanted to identify a new lower threshold for template sequence identity to generate accurate models. We devised an experiment where we binned available templates into groups with 15-19%, 20-24%, 25-29%, and 30-39% sequence identity. We then identified three receptors with at least five templates in each identity group and performed multiple-template homology modeling with each set of five templates. The results, shown in Figure 7, find that overall the TM Region accuracy is unaffected by the use of templates down to 20% sequence identity. The same trend held true for the full model RMSDs. ECL2 was the most sensitive region where accuracy drops sharply when lower identity templates are used. Taken together, we suggest that templates down to 20% identity yield accurate models, particularly within the TM region which is important for ligand recognition.

**Figure 7:**
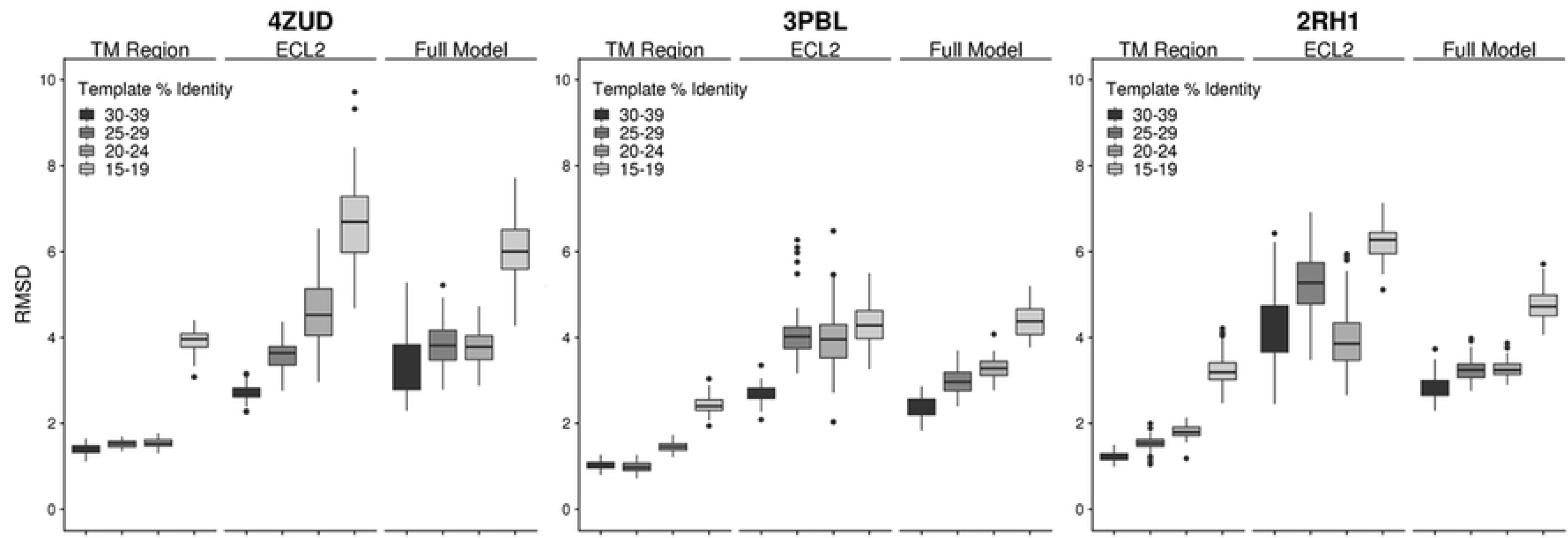
Model Accuracy with Templates over Multiple Sequence Identity Ranges. Three receptors (4ZUD [46], 3PBL [47], and 2RH1 [5]) were identified that had at least 5 templates in each identity range (30-39%, 25-29%, 20-24%, and 15-19%). Using the identified 5 templates for each identity range, 100 models were generated and the RMSD of these models is displayed in box and whisker plots for each region (TM, ECL2, and Full Model).

### Development of Database for All Human Non-Odorant GPCRs

By effectively pushing the lower threshold to 20% sequence identity, we can now predict models for the remaining GPCRs deemed druggable. To this effect, we identified the best templates by identity for the entire set of non-olfactory human GPCRs Figure S2. Out of 397 receptors, 54 have at least one structure determined (14% of the receptor family). This provides 81 receptors with a template with sequence identity above 40%, the previous threshold for accurate modeling. However, the number of receptors with a template between 20 and 40% is 214 (54%). Only 48 receptors (or 12% of the receptor family) remain with sequence identities less than 20% and thus expected lower accuracy in their models. Due to our expectation to be capable of creating accurate structural models over the majority of members of the GPCR family, a model database for all GPCRs without an experimentally determined structure was created using this new method (www.rosettagpcr.org). This is, to the best of our knowledge, the only Rosetta-based GPCR server available which distinguishes it from the many Modeller-based servers. Currently all models are in the inactive state though will likely expand to active state models as more active state structures are determined.

## Discussion

### Blending of Sequence- and Structure-Based Alignment are Critical for Low Identity Template Based Modeling

Inherent to any homology modeling protocol is an alignment between the target and the template sequences. This alignment maps the target sequence onto the template structure in a process called threading [9]. Sequence alignments are necessary for this process and a wide variety of search methods have been generated [27,28]. Each sequence alignment method uses a different algorithm to weight the importance of sequence conservation globally or locally with or without gap penalties. As we learn more about the structures of diverse proteins, it becomes apparent that structure is often more well conserved than sequence. As such, additional algorithms have been generated based on structural alignments and domain fold recognition [29,30]. This latter case is inherent to the family of GPCRs in which the common sequence identity between receptors is around 30% while all receptors share a similar structural domain. Therefore, any method for aligning GPCRs for the purpose of homology modeling should upweight structural alignments over sequence. That is not to say GPCRs lack critical sequence motifs. The NPxxY, CWxP, and DRY motifs as well as numerous proline residues and disulfides are critical for receptor function and should be maintained in their sequence alignment. Of all the alignment methods available, we find that a blended sequence- and structure-based alignment is best for modeling of GPCRs with our method.

One region that is often overlooked when aligning proteins are the loop regions. These loops are highly diverse in sequence as they have evolved over time to recognize many different ligands [36]. However, as more GPCR structures become available, the loop regions are found to adopt similar conformations. As there is a degree of structural conservation, we focused on aligning the loops structurally. This in turn affected how we modeled the loops. By providing a structure-based alignment of the loops in the overall alignment, we allow Rosetta to build the loops simultaneously during the receptor modeling step. This circumvents the need for a secondary loop-closure modeling step as in our previous single-template modeling protocol [22]. This structural alignment and simultaneous loop building contributed significantly to our alignment method outperforming all other alignment methods tested. As shown, almost all methods could accurately align and model the TM region, but only our method performed well modeling the ECL2 region which reduced the full model RMSD. This blend of sequence- and structure-based alignments may prove critical to the modeling of other protein families in which available templates have low sequence identity but high structural conservation. Comparative modeling protocols for antibodies already include sophisticated approaches to categorize and template complementary determining loop regions (CDRs) [41,42].

### Template and Peptide Hybridization are Key Drivers for Accurate Modeling

The Rosetta code for multiple-template comparative modeling (RosettaCM) made two primary changes to the method of homology modeling [23]. The first was the ability to leverage the peptide fragment library derived from structures in the PDB. The peptide fragment library is a set of 3-mers and 9-mers that are mined from the PDB based on the target sequence. These fragments help bias the local geometries of the predicted structures towards structures that are known to exist and are low energy for these short regions [24]. The ability to swap in peptide fragments is helpful for homology modeling, even of single templates, because it allows sampling away from the starting template structure in regions that do not serve well for the target structure. We showed here that threading alone, even for very high identity templates, can be further improved solely by incorporation of peptide fragment hybridization.

The other key change was the use of multiple templates. The original authors of Rosetta’s multiple template homology modeling showed that the use of multiple structures is better than using a single structure [23]. The reasoning is the same as stated above that for a given target, the best available template is still likely to have one or more regions that don’t accurately represent geometries accessible to the target sequence. However, as multiple related templates are used, the likelihood of finding more optimal local structures from combination of the different templates increases. For GPCRs, this has been noted before and many GPCR modeling protocols now use multiple templates. However, their treatment of the multiple templates is distinct than the process used in RosettaGPCR. Often these methods try to pre-identify which regions of the target protein are best captured by each target, and the segments of each template serve as the starting model for energetic minimization and loop rebuilding. In Rosetta, all templates are passed to the hybridization protocol which randomly swaps segments of templates throughout the entire protein. After each modification, the structure is reevaluated and if the score improves, the segment is maintained. As a result, segments that perhaps contained initial lower sequence identity and would have been discarded before the modeling began in other methods, may prove to have better geometries and energies for the target sequence. This selection by energy, not identity, can lead to sampling of more native like conformations for the target sequence. Combined with the peptide fragment insertion, Rosetta can quickly identify native-like conformations distinct from any given template and perform dense sampling around this novel conformation. This is not to suggest that Rosetta will sample conformational changes on the scale of receptor activation (~10 Å in TM 6), but around 1-2 Å of the input templates. While none of this code is new to Rosetta, the application and focus on GPCRs is novel. We present here the method specific for GPCRs and identify the optimal number of templates. Further by optimizing the alignment method and simultaneous loop modeling, we were able to push the previously reported template threshold of a 40% sequence identity minimum down to 20% for accurate modeling.

### Development of the First Rosetta-based GPCR Database

While several GPCR model databases or on-the-fly modeling servers exist, there is as-of-yet one developed within Rosetta. The Robetta server is available for general homology modeling [43], but is not designed for membrane proteins. Further, it uses automatic alignment methods which we showed are not ideal for this class of proteins. Therefore, we decided to generate a database of all non-olfactory human GPCRs (www.rosettagpcr.org). We have confidence in models with templates above 20% sequence identity which accounts for 88% of the receptors. It is important to note that this method was benchmarked on inactive state structures, and therefore, the models in the database are inactive models. These can serve as structures for understanding biochemical data and genetic variations. Further, they can serve as starting points for *in silico* docking campaigns. It has been shown before that docking into inactive structures can work for both identification of agonists and antagonists [44,45]. However, work is ongoing to develop the database further to include active state structures.

### Conclusions

Accurate modeling of GPCRs is a critical technology for understanding the structural basis of ligand recognition and signal transduction for the remaining 350 non-olfactory GPCRs that have not had structures determined. Many of these proteins already have FDA approved drugs targeting them [3], but a deep understanding of the molecular basis of drug intervention is lacking. Further, about a third of these receptors are classified as orphan receptors because the endogenous ligand has not been identified [8]. A structural perspective of the ligand binding pocket may help shed light on this group of receptors. Lastly, it should be noted that this protocol is dependent on novel information. As new structures become available, the template dataset will increase as will the accuracy of the alignment and resulting models. Despite this, in the current format, RosettaGPCR stands as the best modeling protocol for GPCRs. Only 48 receptors remain with sequence identities less than 20% to an available template and thus expected lower accuracy in their models. This subset should be the priority for future experimental structure determination.

## Acknowledgements

This work was conducted in part using the resources of the Advanced Computing Center for Research and Education at Vanderbilt University, Nashville, TN. Funding in the Meiler lab comes from NIH NIGMS R01 GM080403, NIH NIDA R01 DA046138, NIH NIHL R01 HL122010, NIH NIGMS R01 GM129261. BJB was additionally supported by T32 GM07628.

## Supporting Information

**Table S1**. **List of Receptors in Benchmark**. Receptor name and the corresponding PDB ID that was used for accuracy measurements.

**Table S2**. **List of Templates for Each Target Ranked by Sequence Identity**. Yellow highlighted templates were not used for general modeling because they have sequence identities greater than 40%. Bolded templates were used for single-template high identity modeling to compare to previous benchmark.

**Figure S1**. **Alignment of receptor sequences**. The alignment for all 34 receptors is shown using Aline [48]. Identical and highly conserved residues are color-coded for easy identification. Alignment available at www.rosettagpcr.org.

**Figure S2**. **Percent Identify of Best Available Template for Every Non-Odorant Human GPCR**. For each receptor in the human genome, the best template was identified in the PDB. The sequence identity of the best available is plotted. Most templates cross the 20% threshold identified as critical for accurate modeling. The previous threshold of 40% identity is highlighted in red, and the new 20% identity threshold is highlighted in black.

## References

1. Fredriksson R, Lagerström MC, Lundin LG, Schiöth HB. The G-protein-coupled receptors in the human genome form five main families. Phylogenetic analysis, paralogon groups, and fingerprints. Mol Pharmacol. 2003. doi:10.1124/mol.63.6.1256

2. Schiöth HB, Lagerström MC. Structural diversity of g proteincoupled receptors and significance for drug discovery. Nat Rev Drug Discov. 2008. doi:10.1038/nrd2518

3. Hauser AS, Attwood MM, Rask-Andersen M, Schiöth HB, Gloriam DE. Trends in GPCR drug discovery: New agents, targets and indications. Nat Rev Drug Discov. 2017. doi:10.1038/nrd.2017.178

4. Palczewski K, Kumasaka T, Hori T, Behnke CA, Motoshima H, Fox BA, et al. Crystal Structure of Rhodopsin: A G Protein-Coupled Receptor.(Illustration). Science (80-). 2000.

5. Cherezov V, Rosenbaum DM, Hanson MA, Rasmussen SGF, Foon ST, Kobilka TS, et al. High-resolution crystal structure of an engineered human β2-adrenergic G protein-coupled receptor. Science (80-). 2007. doi:10.1126/science.1150577

6. Rasmussen SGF, Choi HJ, Rosenbaum DM, Kobilka TS, Thian FS, Edwards PC, et al. Crystal structure of the human β2 adrenergic G-protein-coupled receptor. Nature. 2007. doi:10.1038/nature06325

7. Bertheleme N, Chae PS, Singh S, Mossakowska D, Hann MM, Smith KJ, et al. Unlocking the secrets of the gatekeeper: Methods for stabilizing and crystallizing GPCRs. Biochimica et Biophysica Acta-Biomembranes. 2013. doi:10.1016/j.bbamem.2013.07.013

8. Sriram K, Insel PA. G protein-coupled receptors as targets for approved drugs: How many targets and how many drugs? Molecular Pharmacology. 2018. doi:10.1124/mol.117.111062

9. Jones DT, Taylort WR, Thornton JM. A new approach to protein fold recognition. Nature. 1992. doi:10.1038/358086a0

10. Šali A, Blundell TL. Comparative Protein Modelling by Satisfaction of Spatial Restraints. J Mol Biol. 1993;234: 779–815. doi:10.1006/jmbi.1993.1626

11. Yang J, Zhang W, He B, Walker SE, Zhang H, Govindarajoo B, et al. Template-based protein structure prediction in CASP11 and retrospect of I-TASSER in the last decade. Proteins. 2016. doi:10.1002/prot.24918

12. Esguerra M, Siretskiy A, Bello X, Sallander J, Gutiérrez-de-Terán H. GPCR-ModSim: A comprehensive web based solution for modeling G-protein coupled receptors. Nucleic Acids Res. 2016. doi:10.1093/nar/gkw403

13. Zhang J, Yang J, Jang R, Zhang Y. GPCR-I-TASSER: A Hybrid Approach to G Protein-Coupled Receptor Structure Modeling and the Application to the Human Genome. Structure. 2015;23: 1538–1549. doi:10.1016/j.str.2015.06.007

14. Miszta P, Pasznik P, Jakowiecki J, Sztyler A, Latek D, Filipek S. GPCRM: A homology modeling web service with triple membrane-fitted quality assessment of GPCR models. Nucleic Acids Res. 2018. doi:10.1093/nar/gky429

15. Worth CL, Kreuchwig F, Tiemann JKS, Kreuchwig A, Ritschel M, Kleinau G, et al. GPCR-SSFE 2.0 - A fragment-based molecular modeling web tool for Class A G-protein coupled receptors. Nucleic Acids Res. 2017. doi:10.1093/nar/gkx399

16. Pándy-Szekeres G, Munk C, Tsonkov TM, Mordalski S, Harpsøe K, Hauser AS, et al. GPCRdb in 2018: Adding GPCR structure models and ligands. Nucleic Acids Res. 2018. doi:10.1093/nar/gkx1109

17. Sandal M, Duy TP, Cona M, Zung H, Carloni P, Musiani F, et al. GOMoDo: A GPCRs Online Modeling and Docking Webserver. PLoS One. 2013. doi:10.1371/journal.pone.0074092

18. Webb B, Sali A. Comparative protein structure modeling using MODELLER. Curr Protoc Bioinforma. 2016. doi:10.1002/cpbi.3

19. Bender BJ, Cisneros A, Duran AM, Finn JA, Fu D, Lokits AD, et al. Protocols for Molecular Modeling with Rosetta3 and RosettaScripts. Biochemistry. 2016;55. doi:10.1021/acs.biochem.6b00444

20. Leaver-Fay A, Tyka M, Lewis SM, Lange OF, Thompson J, Jacak R, et al. Rosetta3: An object-oriented software suite for the simulation and design of macromolecules. Methods in Enzymology. 2011. doi:10.1016/B978-0-12-381270-4.00019-6

21. Kufareva I, Rueda M, Katritch V, Stevens RC, Abagyan R. Status of GPCR Modeling and Docking as Reflected by Community-wide GPCR Dock 2010 Assessment. Structure. 2011;19: 1108–1126. doi:10.1016/J.STR.2011.05.012

22. Nguyen ED, Norn C, Frimurer TM, Meiler J. Assessment and Challenges of Ligand Docking into Comparative Models of G-Protein Coupled Receptors. Seifert R, editor. PLoS One. 2013;8: e67302. doi:10.1371/journal.pone.0067302

23. Song Y, DiMaio F, Wang RY-RYR, Kim D, Miles C, Brunette T, et al. High-resolution comparative modeling with RosettaCM. Structure. 2013;21: 1735–1742. doi:10.1016/j.str.2013.08.005

24. Rohl CA, Strauss CEM, Misura KMS, Baker D. Protein Structure Prediction Using Rosetta. Methods Enzymol. 2004. doi:10.1016/S0076-6879(04)83004-0

25. Larsson P, Wallner B, Lindahl E, Elofsson A. Using multiple templates to improve quality of homology models in automated homology modeling. Protein Sci. 2008. doi:10.1110/ps.073344908

26. Isberg V, Mordalski S, Munk C, Rataj K, Harpsøe K, Hauser AS, et al. GPCRdb: An information system for G protein-coupled receptors. Nucleic Acids Res. 2016. doi:10.1093/nar/gkv1178

27. Sievers F, Higgins DG. Clustal omega, accurate alignment of very large numbers of sequences. Methods Mol Biol. 2014. doi:10.1007/978-1-62703-646-7_6

28. Edgar RC. MUSCLE: A multiple sequence alignment method with reduced time and space complexity. BMC Bioinformatics. 2004. doi:10.1186/1471-2105-5-113

29. Notredame C, Higgins DG, Heringa J. T-coffee: A novel method for fast and accurate multiple sequence alignment. J Mol Biol. 2000. doi:10.1006/jmbi.2000.4042

30. Konagurthu AS, Whisstock JC, Stuckey PJ, Lesk AM. MUSTANG: A multiple structural alignment algorithm. Proteins Struct Funct Genet. 2006. doi:10.1002/prot.20921

31. Viklund H, Elofsson A. OCTOPUS: Improving topology prediction by two-track ANN-based preference scores and an extended topological grammar. Bioinformatics. 2008. doi:10.1093/bioinformatics/btn221

32. Fleishman SJ, Leaver-Fay A, Corn JE, Strauch EM, Khare SD, Koga N, et al. Rosettascripts: A scripting language interface to the Rosetta Macromolecular modeling suite. PLoS One. 2011. doi:10.1371/journal.pone.0020161

33. Ballesteros JA, Weinstein H. Integrated methods for the construction of three-dimensional models and computational probing of structure-function relations in G protein-coupled receptors. Methods Neurosci. 1995;25: 366–428. doi:10.1016/S1043-9471(05)80049-7

34. Isberg V, De Graaf C, Bortolato A, Cherezov V, Katritch V, Marshall FH, et al. Generic GPCR residue numbers - Aligning topology maps while minding the gaps. Trends in Pharmacological Sciences. 2015. doi:10.1016/j.tips.2014.11.001

35. Wheatley M, Wootten D, Conner M, Simms J, Kendrick R, Logan R, et al. Lifting the lid on GPCRs: the role of extracellular loops. Br J Pharmacol. 2012;165: 1688–1703. doi:10.1111/j.1476-5381.2011.01629.x

36. Peeters MC, van Westen GJP, Li Q, IJzerman AP. Importance of the extracellular loops in G protein-coupled receptors for ligand recognition and receptor activation. Trends Pharmacol Sci. 2011;32: 35–42. doi:10.1016/j.tips.2010.10.001

37. Liu H, Kim HR, Deepak RNVK, Wang L, Chung KY, Fan H, et al. Orthosteric and allosteric action of the C5a receptor antagonists. Nat Struct Mol Biol. 2018. doi:10.1038/s41594-018-0067-z

38. Yang Z, Han S, Keller M, Kaiser A, Bender BJ, Bosse M, et al. Structural basis of ligand binding modes at the neuropeptide y Y_1_ receptor. Nature. 2018;556. doi:10.1038/s41586-018-0046-x

39. Cao C, Tan Q, Xu C, He L, Yang L, Zhou Y, et al. Structural basis for signal recognition and transduction by platelet-activating-factor receptor. Nat Struct Mol Biol. 2018. doi:10.1038/s41594-018-0068-y

40. Wang S, Che T, Levit A, Shoichet BK, Wacker D, Roth BL. Structure of the D2 dopamine receptor bound to the atypical antipsychotic drug risperidone. Nature. 2018;555: 269–273. doi:10.1038/nature25758

41. Weitzner BD, Jeliazkov JR, Lyskov S, Marze N, Kuroda D, Frick R, et al. Modeling and docking of antibody structures with Rosetta. Nat Protoc. 2017;12: 401–416. doi:10.1038/nprot.2016.180

42. Finn JA, Koehler Leman J, Willis JR, Cisneros A, Crowe JE, Meiler J. Improving Loop Modeling of the Antibody Complementarity-Determining Region 3 Using Knowledge-Based Restraints. Ho M, editor. PLoS One. 2016;11: e0154811. doi:10.1371/journal.pone.0154811

43. Kim DE, Chivian D, Baker D. Protein structure prediction and analysis using the Robetta server. Nucleic Acids Res. 2004. doi:10.1093/nar/gkh468

44. Kolb P, Rosenbaum DM, Irwin JJ, Fung JJ, Kobilka BK, Shoichet BK. Structure-based discovery of beta2-adrenergic receptor ligands. Proc Natl Acad Sci U S A. 2009;106: 6843–8. doi:10.1073/pnas.0812657106

45. De Graaf C, Rognan D. Selective Structure-Based Virtual Screening for Full and Partial Agonists of the β2 Adrenergic Receptor. J Med Chem. 2008;51: 4978–4985. doi:10.1021/jm800710x

46. Zhang H, Unal H, Desnoyer R, Han GW, Patel N, Katritch V, et al. Structural Basis for Ligand Recognition and Functional Selectivity at Angiotensin Receptor. J Biol Chem. 2015/10/01. 2015;290: 29127–29139. doi:10.1074/jbc.M115.689000

47. Chien EYT, Liu W, Zhao Q, Katritch V, Han GW, Hanson MA, et al. Structure of the human dopamine D3 receptor in complex with a D2/D3 selective antagonist. Science (80-). 2010. doi:10.1126/science.1197410

48. Bond CS, Schüttelkopf AW. ALINE: A WYSIWYG protein-sequence alignment editor for publication-quality alignments. Acta Crystallogr Sect D Biol Crystallogr. 2009. doi:10.1107/S0907444909007835

